# Microbiome stability in wild and rehabilitated insectivorous bats revealed by shotgun metagenomics

**DOI:** 10.64898/2026.02.24.707816

**Authors:** Dongsheng Luo, Alise J. Ponsero, Kate Wright, David J Baker, Andrea Telatin, Colin Townsley, Efstathios S Giotis

## Abstract

**Background:** Wildlife rehabilitation can influence host-associated microbiota, yet little is known about how the gut microbiome of insectivorous bats responds to rehabilitation during temporary managed care. This study applied shotgun metagenomics to evaluate the impact of temporary managed care on the gut microbiome of wild and rehabilitated bats in Yorkshire, UK.

**Results:** We analysed 25 faecal metagenomes from *Myotis daubentonii, Pipistrellus pipistrellus, Nyctalus noctula* and *N. leisleri*, including wild baseline bats and bats sampled during temporary managed care (1-49 days in rehabilitation). Microbial communities clustered strongly by host species and roost location, but not by rehabilitation status. Bacterial alpha diversity did not differ between wild bats, and bats in care (H = 2.30, *p* = 0.32). Archaeal communities were highly uniform across samples, showing far lower interindividual variation than bacterial communities (12.2% vs. 41.8% coefficient of variation). Rehabilitated bats showed increased relative abundance of *Yersiniaceae* and *Lactobacillaceae*, while environmental families such as *Pseudomonadaceae* and *Erwiniaceae* decreased, indicating modest but non-disruptive changes consistent with a controlled diet and reduced environmental exposure.

**Conclusions:** Across temporary managed care, the core gut microbiome of insectivorous bats remained stable, demonstrating notable microbial resilience. These findings provide an important baseline for monitoring microbiome health in wildlife rehabilitation and supporting post-release conservation programmes in the UK and beyond.

## Background

Bats are ecologically vital and legally protected species in the United Kingdom, contributing to insect population control, pollination, and broader ecosystem balance [1]. However, many species are increasingly threatened by habitat loss, food scarcity, and anthropogenic pressures that result in injury, illness, or displacement [2, 3]. The gut microbiome is central to animal health, mediating nutrient absorption, immune regulation, and resistance to pathogens [4]. In mammals, captivity and dietary change can substantially alter microbial diversity and composition, often reducing ecological complexity and enriching opportunistic taxa [5-8].

Despite the importance of rehabilitation in conservation programmes, the effects of short-term rehabilitation and controlled diets on the gut microbiome of insectivorous bats remain unexplored [9-12]. This gap is particularly relevant given the limited understanding of the role of the bat gut microbiome in pathogen tolerance, and that certain bat species harbour a wide range of viruses, including coronaviruses, rhabdoviruses, paramyxoviruses, and influenza-like viruses, reflecting their long evolutionary associations with diverse viral families [13-17]. Rehabilitation feeding typically relies on mealworms and other commercially reared insects, providing a simplified diet that may alter microbial composition relative to that of wild prey [18].

In the UK, no previous studies have examined gut microbiome responses of insectivorous bats during rehabilitation. This represents a critical gap, as temporary captivity is common in bat conservation practice. Understanding whether short-term rehabilitation alters microbial composition is essential for ensuring welfare standards, maintaining ecological function, and supporting post-release adaptation [12]. In this study, we used shotgun metagenomic sequencing to investigate the gut microbiome of four insectivorous bat species, *Myotis daubentonii, Pipistrellus pipistrellus, Nyctalus noctula*, and *N. leisleri*, from wild roosts and rehabilitation facilities in Yorkshire, UK. We compared microbial diversity and composition between wild bats and bats sampled during temporary managed care and assessed whether microbiome patterns varied with time in care (1–49 days).

## Methods

### Study design and sampling

Thirty-three faecal DNA extracts were initially collected from insectivorous bats across Yorkshire, UK, including individuals sampled from wild roosts and rehabilitation facilities coordinated by the West Yorkshire Bat Group (Supplementary Table S1). Sampling was non-invasive, with fresh faecal pellets collected from clean surfaces beneath roosts or within holding enclosures during rehabilitation. Following quality filtering, 25 samples were retained: 14 from wild baseline bats and 11 from bats sampled during rehabilitation (Supplementary Table S2). Rehabilitation duration was determined from admission and sampling records provided by bat carers registered with the Bat Conservation Trust. Bats in managed care were sampled between 1 and 49 days after admission. Most samples (C01–C03 and C07–C11) were collected within 7 days of care, whereas three samples (C04–C06) were collected after extended care of 42–49 days. Samples labelled W represent wild baseline bats sampled directly from roosts. Species-level identity was resolved for 18 of the 25 samples, representing four confirmed species: *Myotis daubentonii, Pipistrellus pipistrellus, Nyctalus noctula* and *Nyctalus leisleri*.

### DNA sequencing and processing

Metagenomic libraries were prepared using standard Illumina protocols and sequenced on a NovaSeq X Plus platform (25B flow cell) at Quadram Institute to generate paired-end 150 bp reads. Raw reads were quality-trimmed with TrimGalore (v0.6.10), to remove low-quality bases and adapter sequences. Human-associated contamination was filtered using Hostile (v2.0.0) with the human-t2t-hla reference [19]. To ensure sufficient sequencing depth for reliable profiling analysis, a threshold of 1 million reads post-QC was applied. Sequencing data have been deposited in the European Nucleotide Archive under accession number PRJEB103765.

### Metagenomic analysis

#### Host identification

Bat host species were identified using a combined assembly- and read-mapping approach. Quality-controlled reads were assembled with MEGAHIT v1.2.9 [20], and contigs were classified as organellar or non-organellar using Tiara v1.0.3 [21]. Organellar contigs were queried against the NCBI nucleotide database (nt v5) using BLASTn v2.10, retaining high-confidence hits (alignment length ≥500 bp, e-value ≤1×10^−10^, percent identity ≥95%). Top-scoring mitochondrial matches were used for species-level identification. To complement this and account for low host DNA content, reads were also mapped to complete bat reference genomes using Bowtie2 v2.5.4, and the reference genome with the highest mapping proportion was used to assign host genus or species identity. Samples with conflicting results or minimal host DNA signal were flagged as “unclear” for host identity. Mapping proportions were visualised as a z-score–normalised heatmap with hierarchical clustering (Euclidean distance, complete linkage) using pheatmap v1.0.12.

#### Prokaryotic profiling

Quality-controlled reads were taxonomically classified using Kraken2 v2.1.3 against the k2_core_nt_20250609 database. Alpha diversity (Shannon index) and beta diversity (PCA) were calculated in R v4.3.3 using the microbiome and phyloseq packages. Statistical comparisons of alpha diversity were performed using Kruskal– Wallis tests followed by pairwise Wilcoxon rank-sum tests where applicable.

#### Eukaryotic profiling

Quality-controlled reads were assembled with MEGAHIT v1.2.9 [20], and contigs were classified using Tiara v1.0.3 [21]. Organellar contigs were annotated by BLASTn against the NCBI nt database (v5) using the same filtering criteria (alignment length ≥500 bp, e-value ≤1×10^−10^, percent identity ≥95%), retaining the top-scoring hit for taxonomic assignment.

#### Viral profiling

Viral sequences were identified from assembled contigs using geNomad v1.8.0 [22]. Viral sequence quality was assessed with CheckV v1.0.3 (database v1.5) [23], retaining sequences classified as medium, high, or complete, or >1 kb in length. A total of 28,371 sequences were clustered into vOTUs (95% identity over 85% of the longest sequence) using aniclust and anicalc (CheckV utilities), yielding 22,730 vOTUs. Quality-controlled reads were mapped to representative vOTU sequences using bwa-mem as implemented in coverM v0.7.0, with a minimum coverage of 75% and read identity of 90%, and abundances were normalised as TPM using coverM v0.7.0 [24].

## Results

After quality control and read-depth filtering (>1 × 10^6^ reads), 25 faecal metagenomes were retained for analysis, comprising 14 wild bat samples from natural roosts and 11 bat samples taken during rehabilitation. Host species identity was determined through mitochondrial genome assembly and reference mapping (Figure 1A, Supplementary Table S2). Species-level identification was achieved for 19 samples: 6 *Myotis daubentonii*, 10 *Pipistrellus pipistrellus*, 1 *Nyctalus leisleri*, and 2 *Nyctalus noctula*. The remaining 6 samples were classified as uncertain host.

**Figure 1.**
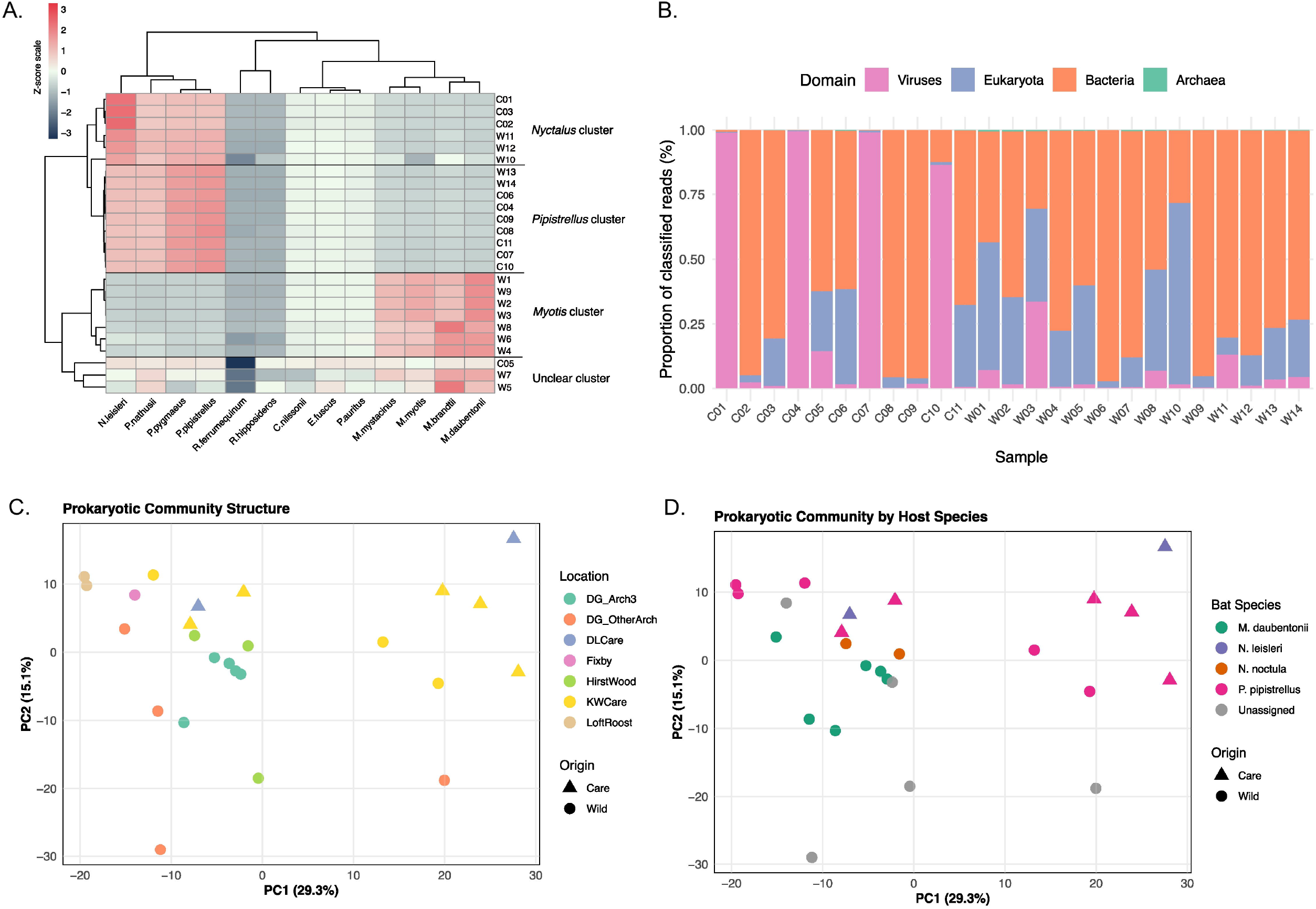
Overview of microbiome composition across wild and rehabilitated bats. (A) Heatmap of the proportion of quality-controlled reads from each fecal sample mapping to bat reference genomes. Values are z-score normalized by sample (rows) to highlight relative mapping preferences. Samples and reference genomes (columns) are hierarchically clustered using Euclidean distance and complete linkage. Samples marked as “unclear” show no distinct clustering pattern to any reference genome cluster, indicating insufficient host DNA content for genus assignment. (B) Domain-level composition of classified reads showing the relative proportions of Bacteria, Eukaryota, Viruses, and Archaea. (C) Principal component analysis (PCA) of prokaryotic community composition coloured by sampling location and shaped by origin (wild vs. care). (D) PCA coloured by host species and shaped by origin (wild vs. care). Both PCA was performed on centered log-ratio (CLR) transformed abundance data representing Aitchison distance.

Prokaryotic sequences dominated the classified fraction of reads (median = 74.7%), followed by eukaryotic (17.3%) and viral sequences (1.6%) (Figure 1B, Supplementary Figure 1). Principal component analysis (PCA) of centred log-ratio-transformed family-level abundances revealed clear clustering by host species and sampling location, but no clear distinction between wild and in-care bats (Figure 1C-D). These factors were confounded in the study design, as species composition differed across sampling locations (e.g., all *M. daubentonii* originated from a specific location: Dowley Gap arches, Supplementary Table S1). Nonetheless, the absence of separation by rehabilitation status indicates that host identity and roost environment exerted stronger effects on microbiome structure than captivity. Bacterial alpha diversity (Shannon index 2.5-3.3) did not differ significantly during the managed care length (Kruskal-Wallis H = 2.30, p = 0.32; Supplementary Figure 1C). However, bacterial community composition showed modest shifts at the family level when comparing wild bats to those in rehabilitation (Figure 2A-B). Mean relative abundance of *Lactobacillaceae* increased from 4.5% in bats in care for less that 4 days to 31.9% in bats in care for a longer period, and *Yersiniaceae* increased from 0.85% to 21.0%. Conversely, families commonly associated with environmental exposure declined: *Erwiniaceae* decreased from 19.3% to 1.2%, and *Pseudomonadaceae* from 9.3% to 0.4% (Figure 2B). The enrichment of *Lactobacillaceae* may reflect acquisition of bacteria from the mealworm (*Tenebrio molitor*) gut microbiome, or an impact of the milk powder supplementation. These compositional shifts were not consistent across individuals and modest in magnitude, suggesting adaptation to a standardised captive diet without broad community disruption.

**Figure 2.**
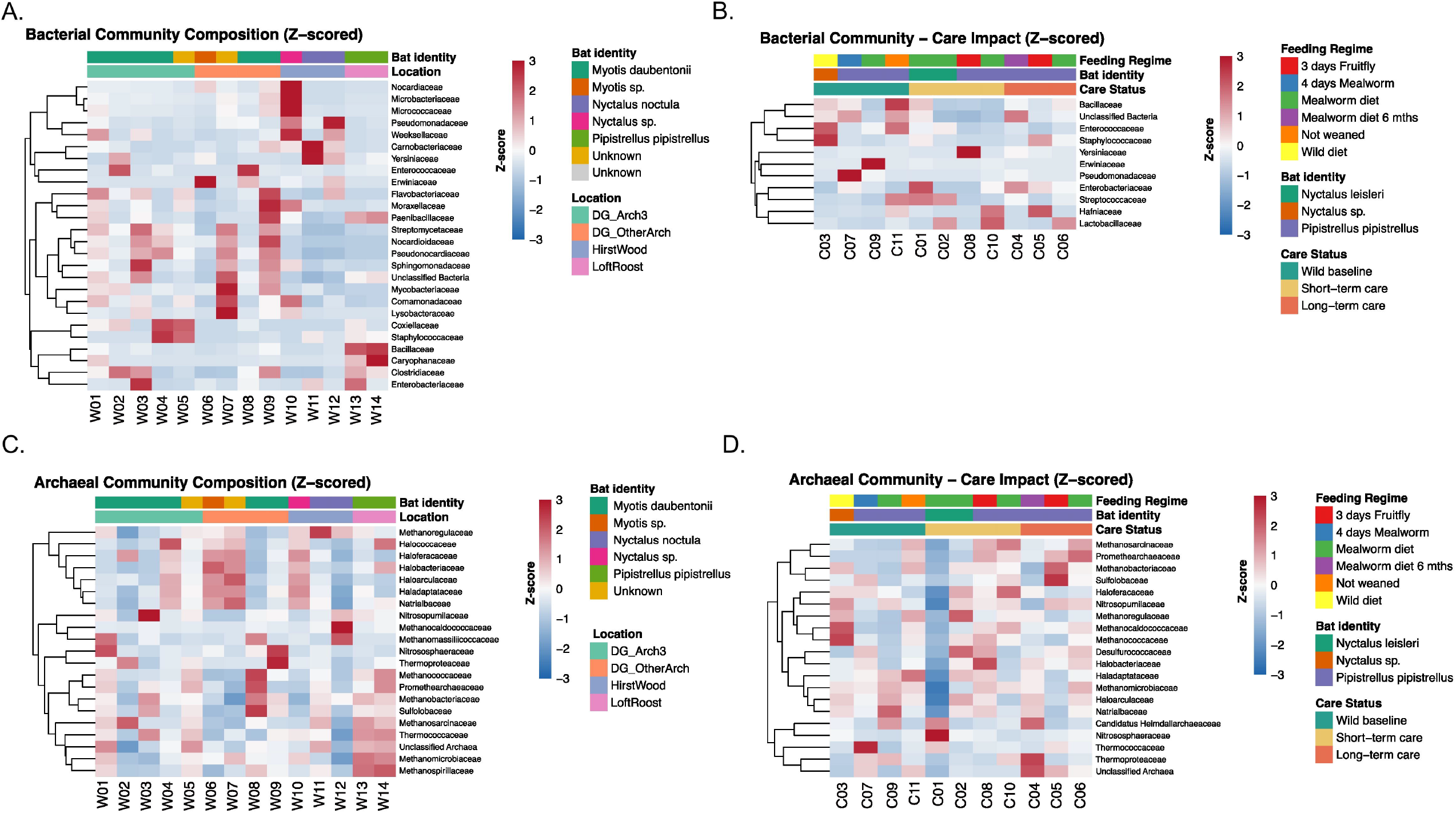
Bacterial and archaeal community composition in wild and managed-care bats. Z-scored relative abundances of dominant prokaryotic families (detected in >1% in ≥3 samples) across wild samples (A – Bacteria; C-Archaea) or bats in care (B – Bacteria; D - Archaea), annotated by host species and sampling location and feeding.

In contrast to the modest compositional shifts observed in bacterial communities, archaeal communities were highly uniform across all samples, dominated by *Methanobacteriaceae* and *Methanomassiliicoccaceae*. Archaeal family-level profiles showed minimal inter-individual variation (12.2% coefficient of variation) compared to bacterial communities (41.8% coefficient of variation), and no clear separation between wild bats and bats in rehabilitation (Figure 2C-D). Archaeal diversity did not differ significantly according to the rehabilitation period length (Kruskal-Wallis H = 0.66, p = 0.72, supplemental figure 1C).

Fungal DNA was detected sporadically in 13 of 25 samples (52%), representing 23 genera dominated by *Penicillium, Mucor*, and *Debaryomyces*. Fungal reads were more frequent in wild roost samples (60%) than in rehabilitation facilities (38%), consistent with environmental or dietary origins rather than established gut colonisation. Parasitic DNA was identified in two wild bats: *Plagiorchis vespertilionis* and *Eimeria jerfinica*, both recognised bat parasites, and *Gordius albopunctatus*, a nematomorph likely derived from infected arthropod prey.

Arthropod mitochondrial sequences revealed clear differences in prey composition between wild and captive bats (Figure 3B). Wild individuals consumed a variety of aquatic and semi-aquatic insects, including mayflies (*Caenis luctuosa, Ephemera danica*), caddisflies (*Hydropsyche contubernalis*), and crane flies (*Tipula helvola*), consistent with foraging over water. In contrast, bats in rehabilitation fed almost exclusively on commercial feeder insects (*Tenebrio molitor*), validating dietary records provided by carers.

**Figure 3.**
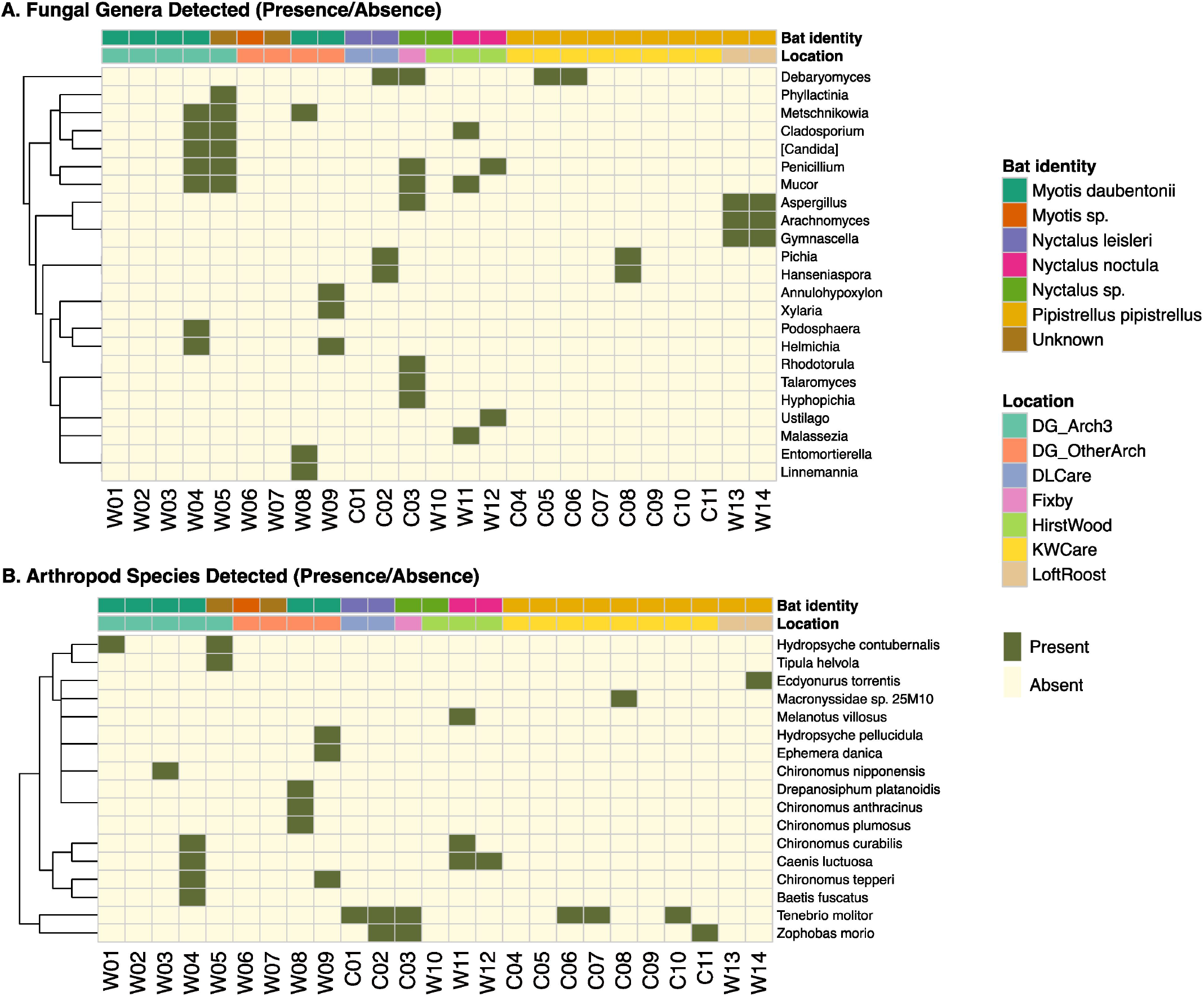
Fungal and Arthropods sequences detected in wild and rehabilitated bats. Heatmap showing the distribution of (A) fungal genera and (B) arthropod genera across bat faecal samples based on organellar BLAST analysis. Each cell represents the presence (green) or absence (cream) of a fungal/arthropod genus in a given sample. Samples (columns) are ordered by collection location (top annotation bar) with bat host species indicated (second bar). Genera (rows) are hierarchically clustered using Euclidean distance to group taxa with similar distribution patterns.

The assembled viral catalogue comprised 22,729 viral operational taxonomic units (vOTUs), dominated by bacteriophages of the realm *Duplodnaviria* (≈90%), followed by *Varidnaviria, Monodnaviria*, and *Riboviria* (Supplementary Figure 2). Putative mammalian-associated viruses were identified by BLAST homology to known viral families and filtered for coverage >75% and mapping identity >90%, yielding 196 candidate vOTUs distributed across families including *Herpesviridae, Parvoviridae*, and *Poxviridae* (Supplementary Table 3). The majority of abundant viral sequences corresponded to dietary or environmental sources, including *Bee densovirus 2* in wild bats and *Zophobas morio black wasting virus* in captive bats, suggesting the presence of a closely related virus infecting an alternative worm host. High-coverage sequences included bat parvovirus 3 in *M. daubentonii* samples and a divergent papillomavirus in *N. noctula* (>1000× coverage), which may represent a novel bat-associated lineage pending further validation (Supplementary Table 4). No sequences related to known zoonotic viruses were detected.

## Discussion

Our study demonstrates that the gut microbiome of insectivorous bats is highly individual specific and remains stable during temporary managed care lasting 1-49 days (including three bats sampled at 42–49 days), despite dietary and environmental changes. Consistent with this, we detected no significant differences in microbial diversity or overall community structure between wild and rehabilitated bats. This contrasts with most mammalian systems, where captivity typically leads to reduced microbial diversity and enrichment of opportunistic or host-associated taxa [5, 25]. One possible explanation is the use of whole-insect diets during rehabilitation, which more closely resemble natural prey and may be less disruptive than the processed or plant-based diets often provided to other captive mammals. In this study’s bats, bacterial α- diversity, archaeal composition, and broader community profiles were preserved, indicating a high degree of microbial resilience. Although sample availability was naturally limited in a rehabilitation context, the consistent patterns across individuals and care durations suggest that this stability is biologically meaningful.

This resilience is also consistent with ecological and evolutionary traits unique to bats. Many species routinely experience fluctuating diets, variable foraging conditions, and rapid physiological shifts associated with heterothermy, which may select for flexible and disturbance-tolerant microbial communities [26, 27]. The clear clustering of samples by host species rather than captivity status supports this idea and reflects a signature of phylosymbiosis, where microbiome composition tracks host evolutionary relationships [8, 26]. This phylogenetic signal suggests long-term co-adaptation between bats and their gut symbionts, which might be helping buffer the microbiome against short-term anthropogenic disturbance.

Microbiome stability in these bats may also reflect general functional redundancy within gut microbial communities, a pattern observed across many vertebrates where core metabolic processes remain conserved even when taxonomic composition shifts [28]. Future metagenomic and metatranscriptomic analyses could confirm whether nutrient-processing and immune-related functions persist unchanged in rehabilitated bats. Together, these findings suggest the presence of a resilient prokaryotic core that maintains overall community structure and diversity despite taxonomic shifts in response to dietary change and captivity.

Dietary DNA analysis revealed expected differences between wild and captive bats, reflecting natural foraging versus standardised feeding in rehabilitation. Wild individuals consumed a diverse range of aquatic and woodland insects, whereas captive bats were fed primarily mealworms, with occasional supplementation. Correspondingly, environmentally associated bacterial families were reduced in captivity, consistent with a simplified prey spectrum [28]. These changes occurred without loss of overall bacterial diversity or alteration of archaeal community structure, supporting an adaptive microbial response rather than microbiome disruption. Dietary enrichment during rehabilitation may therefore help maintain microbial diversity and metabolic flexibility.

Beyond bacteria and archaea, fungal and viral reads were detected but were largely environmental or dietary in origin. No sequences related to known zoonotic viruses were found. A divergent papillomavirus signal was identified in one *N. noctula* sample, but additional targeted sequencing would be needed to confirm whether it represents a true bat-associated virus.

From a conservation standpoint, the results highlight that temporary rehabilitation lasting up to 49 days, when conducted with minimal stress and dietary care, does not disrupt the microbiome integrity of bats. Maintaining this microbial stability is crucial because gut microbes contribute to nutrient assimilation, immune function, and post-release adaptation [6]. Microbiome profiling can therefore serve as an early indicator of rehabilitation success and welfare.

Several limitations of this study should be acknowledged. First, the cross-sectional sampling design means that each bat was sampled at a single time point, precluding longitudinal tracking of microbiome trajectories within individuals over the course of rehabilitation. Second, host species identity, roost location, feeding diet and rehabilitation status are partially confounded in this dataset which limits the ability to disentangle the relative contributions of these factors to microbiome variation. Finally, as sampling was necessarily opportunistic within a rehabilitation context, group sizes were limited and uneven, which may constrain the detection of subtle microbiome differences and warrants cautious interpretation of small effect sizes.

Future directions should include longitudinal tracking of individual bats from capture through release, integrating metagenomic, metabolomic, and physiological data. Linking microbial shifts to health markers, such as body condition, immune gene expression, and thermoregulatory capacity will clarify functional consequences of captivity and reintroduction. Expanding comparative datasets across bat guilds (insectivorous, frugivorous, nectarivorous) will also test whether the resilience observed here is a broader chiropteran trait or specific to temperate insectivores.

## Conclusions

Shotgun metagenomic sequencing of individual bat droppings provides a powerful, non-invasive framework for integrated assessment of host identity, diet, and microbial health. Despite simplified diets and captive conditions, insectivorous bats retained diverse and functionally redundant microbial consortia, demonstrating exceptional microbiome resilience. This contrasts with the pronounced microbiome alterations reported in many captive mammals and suggests that bats might be more tolerant to environmental and dietary variation. By revealing stability across bacterial and archaeal domains, our study supports the use of metagenomic profiling as a benchmark for bat welfare, and rehabilitation standards. Future integration of functional metagenomics and longitudinal monitoring will strengthen the link between microbial resilience and ecological fitness, ensuring that rehabilitation efforts sustain not only the host but also its symbiotic microbiota.

## Declarations

### Ethics approval and consent to participate

Guano samples were collected non-invasively by bat carers registered with the Bat Conservation Trust, UK. DNA extraction was performed at the University of Essex under institutional biosafety and ethical approval (Ref: UoE-BIO-LS-25-001 and ETH2526-0296).

## Consent for publication

Not applicable.

## Availability of data and materials

Sequencing data have been deposited in the European Nucleotide Archive under accession number PRJEB103765.

## Authors’ contributions

DL performed DNA isolation and quality control. KW and CT collected samples. ESG and CT conceptualised the study and contributed to its design. DB performed the sequencing of the dataset. AP and AT conducted the bioinformatic analyses. ESG wrote the manuscript with input from all authors. All authors read and approved the final version.

## Acknowledgements

We thank Dan Lindley and Paul Redmond for assistance with the collection of wild samples, and Alan Brailsford for coordinating work between the Quadram Institute and the University of Essex. We also thank University of Essex personnel for sample extraction support and the West Yorkshire Bat Group for assistance with field logistics.

