## Supplementary material for "Microbiome stability in wild and rehabilitated insectivorous bats revealed by shotgun metagenomics"

### Supplementary Table S1. Overview of bat faecal samples by location and rehabilitation status

This table summarises faecal samples collected from wild bat roosts and rehabilitation centres in Yorkshire, UK. Samples are grouped by location/site, care status, and rehabilitation duration. For each site, the manuscript IDs of associated samples, total number of samples, and host species detected (based on mitochondrial confirmation; see Supplementary Table S2 and read mapping) are reported.

| Location / Site | Area | Care status | Rehabilitation duration | Manuscript IDs | No. samples | Species detected |
|---|---|---|---|---|---|---|
| Dowley Gap | Bradford (West Yorkshire) | Wild roost | Baseline | W1, W2, W3, W4, W5 | 5 | *Myotis daubentonii* |
| Dowley Gap 2 | Bradford (West Yorkshire) | Wild roost | Baseline | W6, W7, W8, W9 | 4 | *Myotis daubentonii* |
| Hirst Wood | Bradford (West Yorkshire) | Wild roost | Baseline | W10, W11, W12 | 3 | *Pipistrellus pipistrellus*, *Nyctalus noctula* |
| Loft Roost | Keighley (West Yorkshire) | Wild roost | Baseline | W13, W14 | 2 | *Pipistrellus pipistrellus* |
| DL Care facility | Kirklees (West Yorkshire) | Rehabilitation | ≤7 days | C01, C02 | 2 | *Myotis daubentonii*, *Nyctalus leisleri* |
| Fixby | Huddersfield area | Rehabilitation | ≤7 days | C03 | 1 | *Uncertain* |
| KW Care facility | North Yorkshire | Rehabilitation | 1-49 days | C04, C05, C06, C07, C08, C09, C10, C11 | 8 | *Pipistrellus pipistrellus* |

### Supplementary Table S2. Host species identification from mitochondrial read mapping

This table provides species-level host identification for all metagenomic faecal samples included in the study. For each sample, the top mitochondrial BLASTn hit (SSEQID), corresponding species, percent identity, and alignment length are shown. Species assignments were based on the highest-identity mitochondrial match (>98% identity and >500 bp alignment length).

| Manuscript ID | Sample Name | Care Status | Rehabilitation duration | Location | SSEQID | Species Hit | Percent Identity | Alignment Length |
|---|---|---|---|---|---|---|---|---|
| **C01** | Care_DL_S17 | Care | ≤7 days | DL care facility | | | | |
| **C02** | Care_DL_S18 | Care | ≤7 days | DL care facility | OZ183647.2 | *Nyctalus leisleri* | 99,81 | 15757 |
| **C03** | Care_Fixby_S16 | Care | ≤7 days | Fixby | | | | |
| **C04** | Care_KW_S20 | Care | 42 days | KW care facility | LR862378.1 | *Pipistrellus pipistrellus* | 99,929 | 15558 |
| **C05** | Care_KW_S21 | Care | 45 days | KW care facility | LR862378.1 | *Pipistrellus pipistrellus* | 99,979 | 4807 |
| **C06** | Care_KW_S22 | Care | 49 days | KW care facility | LR862378.1 | *Pipistrellus pipistrellus* | 99,929 | 15558 |
| **C07** | Care_KW_S23 | Care | 4 days | KW care facility | LR862378.1 | *Pipistrellus pipistrellus* | 99,929 | 15559 |
| **C08** | Care_KW_S24 | Care | 7 days | KW care facility | LR862378.1 | *Pipistrellus pipistrellus* | 99,929 | 15559 |
| **C09** | Care_KW_S26 | Care | 1 day | KW care facility | LR862378.1 | *Pipistrellus pipistrellus* | 99,93 | 15640 |
| **C10** | Care_KW_S28 | Care | 6 days | KW care facility | LR862378.1 | *Pipistrellus pipistrellus* | 99,93 | 15640 |
| **C11** | Care_KW_S30 | Care | 1 day | KW care facility | LR862378.1 | *Pipistrellus pipistrellus* | 99,852 | 15580 |
| **W1** | Wild_DG_S1 | Wild | Baseline | Dowley Gap | MN122860.1 | *Myotis daubentonii* | 99.854 | 15757 |
| **W2** | Wild_DG_S2 | Wild | Baseline | Dowley Gap | MN122860.1 | *Myotis daubentonii* | 99.888 | 15134 |
| **W3** | Wild_DG_S3 | Wild | Baseline | Dowley Gap | OY725382.2 | *Myotis daubentonii* | 99.749 | 15919 |
| **W4** | Wild_DG_S5 | Wild | Baseline | Dowley Gap | MN122860.1 | *Myotis daubentonii* | 99.754 | 15838 |
| **W5** | Wild_DG_S6 | Wild | Baseline | Dowley Gap | | | | |
| **W6** | Wild_DG2_S10 | Wild | Baseline | Dowley Gap 2 | | | | |
| **W7** | Wild_DG2_S12 | Wild | Baseline | Dowley Gap 2 | | | | |
| **W8** | Wild_DG2_S7 | Wild | Baseline | Dowley Gap 2 | MN122860.1 | *Myotis daubentonii* | 99.874 | 15822 |
| **W9** | Wild_DG2_S8 | Wild | Baseline | Dowley Gap 2 | MN122860.1 | *Myotis daubentonii* | 99.911 | 15702 |
| **W10** | Wild_HirstWood_S 13 | Wild | Baseline | Hirst Wood | | | | |
| **W11** | Wild_HirstWood_S 14 | Wild | Baseline | Hirst Wood | MN122907.1 | *Nyctalus noctula* | 99.815 | 15648 |
| **W12** | Wild_HirstWood_S 15 | Wild | Baseline | Hirst Wood | MN122907.1 | *Nyctalus noctula* | 99.802 | 15648 |
| **W13** | Wild_LoftRoost_S 31 | Wild | Baseline | Loft Roost | LR862378.1 | *Pipistrellus pipistrellus* | 99.955 | 15558 |
| **W14** | Wild_LoftRoost_S 32 | Wild | Baseline | Loft Roost | LR862378.1 | *Pipistrellus pipistrellus* | 99.904 | 15558 |

**Supplementary Figure 1. Taxonomic classification efficiency of metagenomic reads in wild and rehabilitated bats**

A) Total number of classified and unclassified reads per sample following Kraken2 taxonomic assignment. Bars represent individual metagenomes (n = 26), with classified reads shown in turquoise and unclassified reads in red. Samples include both wild bats and individuals in rehabilitation.
B) Proportion of unclassified reads (%) in rehabilitated (Care) and wild bats. Wild samples showed a significantly higher proportion of unclassified reads than rehabilitated bats (Wilcoxon rank-sum test, $p$ = 0.014), consistent with increased dietary and environmental diversity in wild conditions.
C) Alpha-diversity (Shannon index at the species level) in the bats in rehabilitation grouped by their stay length in care

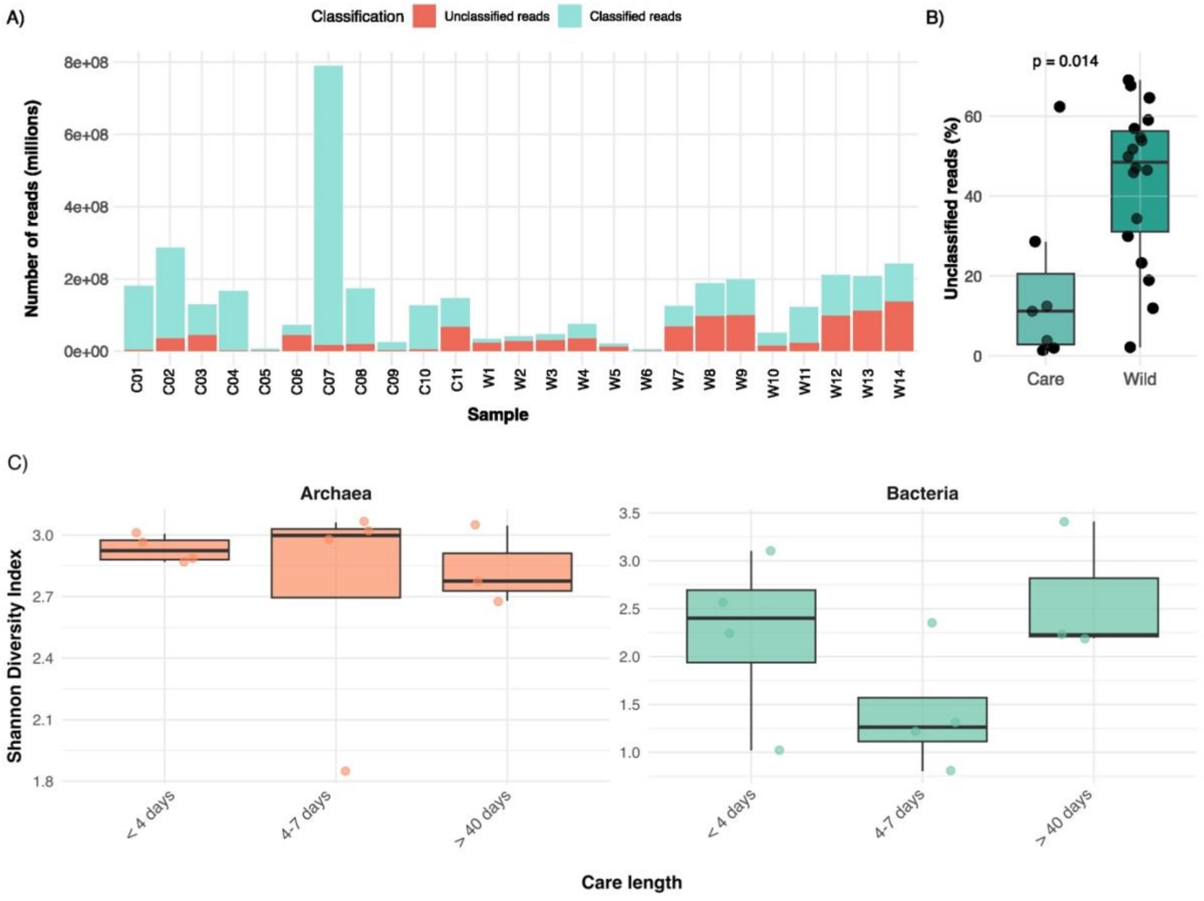

**Supplementary Figure 2. Composition of the bat virome across all metagenomic samples.** Distribution of viral operational taxonomic units (vOTUs) detected across all 26 bat metagenomes grouped by viral realm. The virome was dominated by bacteriophages of the realm Duplodnaviria (20,565 vOTUs; 90.5%), followed by smaller contributions from Varidnaviria, Monodnaviria, and Riboviria. A minority of vOTUs were unclassified or belonged to unranked viral groups. The predominance of phage-associated realms reflects the bacterial origin of most viral sequences within gut metagenomes.

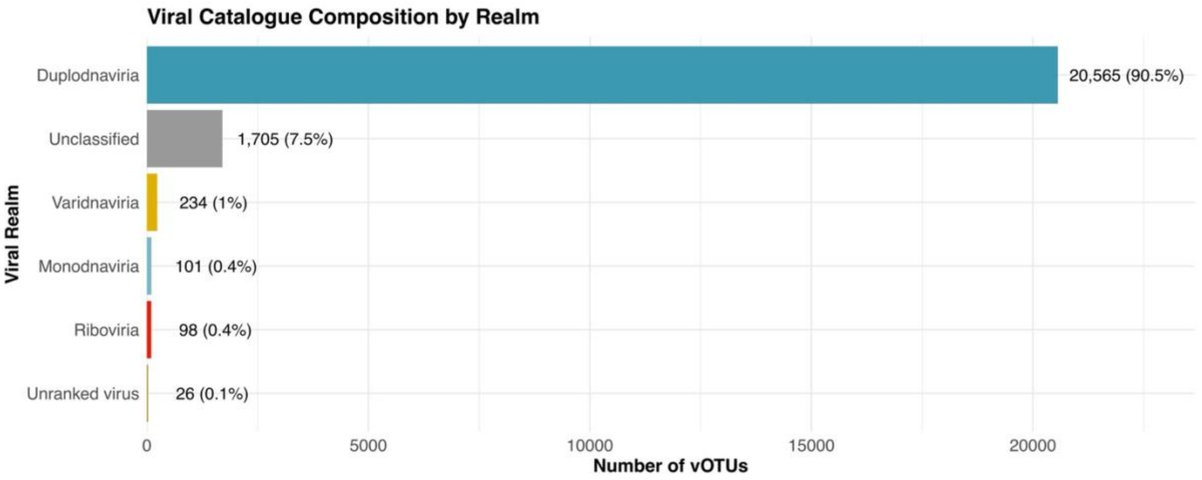

**Supplementary Table 3. Viral families detected across all bat metagenomes**

Summary of viral operational taxonomic units (vOTUs) assigned to each viral family across the 26 metagenomic samples. Viral families are grouped by ecological or host association: mammalian viruses, invertebrate viruses, fungal viruses, environmental viruses, and arthropod/plant viruses. For each family, vOTUs and their percentage contribution to the total viral catalogue are reported. These values reflect the expected predominance of dietary, environmental, and bacteriophage-derived sequences in gut metagenomes.

| Viral Family | Number of vOTUs | % of Catalogue |
|---|---|---|
| **Mammalian virus** | 196 | 0,87 |
| **Herpesviridae** | 43 | 0,19 |
| **Retroviridae** | 41 | 0,18 |
| **Poxviridae** | 39 | 0,17 |
| **Parvoviridae** | 36 | 0,16 |
| **Papillomaviridae** | 16 | 0,07 |
| **Adenoviridae** | 13 | 0,06 |
| **Polyomaviridae** | 8 | 0,04 |
| **Invertebrate virus** | 92 | 0,4 |
| **Adintoviridae** | 39 | 0,17 |
| **Iridoviridae** | 30 | 0,13 |
| **Baculoviridae** | 23 | 0,1 |
| **Fungal virus** | 25 | 0,11 |
| **Totiviridae** | 18 | 0,08 |
| **Partitiviridae** | 7 | 0,03 |
| **Environmental virus** | 88 | 0,38 |
| **Mimiviridae** | 51 | 0,22 |
| **Phycodnaviridae** | 30 | 0,13 |
| **Lavidaviridae** | 7 | 0,03 |
| **Arthropod/Plant virus** | 8 | 0,04 |
| **Phenuiviridae** | 8 | 0,04 |

**Supplementary Table 4. High-coverage viral sequences detected in individual samples.** Viral operational taxonomic units (vOTUs) achieving notable sequencing coverage (>50×) in individual bat samples. Columns include sample identifier, confirmed bat species, inferred viral realm, sequencing coverage, and closest reference match. High-coverage viral hits included Parvoviridae-like sequences (Parvovirus bat6352) in several wild *Myotis daubentonii* and *Nyctalus noctula* samples, and two Duplodnaviria hits corresponding to endogenous bat chromosomal sequences in *Pipistrellus pipistrellus*.

| ID | Sample_short | Bat | Realm | Coverage | Closest_hit |
|---|---|---|---|---|---|
| C10 | Care_KW_S28 | Pipistrellus pipistrellus | Duplodnaviria | 100 | bat chromosome |
| C04 | Care_KW_S20 | Pipistrellus pipistrellus | Duplodnaviria | 82 | bat chromosome |
| C05 | Care_KW_S21 | Pipistrellus pipistrellus | Monodnaviria | 4108 | |
| W12 | Wild_Hirst Wood_S15 | Nyctalus noctula | Monodnaviria | 1405 | Parvovirus bat6352 |
| W2 | Wild_DG_S2 | Myotis daubentonii | Monodnaviria | 94 | Parvovirus bat6352 |
| W3 | Wild_DG_S3 | Myotis daubentonii | Monodnaviria | 53 | Parvovirus bat6352 |
| W12 | Wild_Hirst Wood_S15 | Nyctalus noctula | Monodnaviria | 12436 | |
| W3 | Wild_DG_S3 | Myotis daubentonii | Monodnaviria | 347 | Parvovirus bat6352 |
| W12 | Wild_Hirst Wood_S15 | Nyctalus noctula | Monodnaviria | 285 | Parvovirus bat6352 |
| W2 | Wild_DG_S2 | Myotis daubentonii | Monodnaviria | 58 | Parvovirus bat6352 |
| W2 | Wild_DG_S2 | Myotis daubentonii | Monodnaviria | 92 | |
| C05 | Care_KW_S21 | Pipistrellus pipistrellus | Monodnaviria | 300 | Protoparvovirus chiropteran2 |
| C04 | Care_KW_S20 | Pipistrellus pipistrellus | Monodnaviria | 54 | Protoparvovirus chiropteran2 |
| C01 | Care_DL_S17 | Nyctalus leisleri | Monodnaviria | 8306177 | Zophobas morio black wasting virus |
| C04 | Care_KW_S20 | Pipistrellus pipistrellus | Monodnaviria | 7439284 | Zophobas morio black wasting virus |
| C10 | Care_KW_S28 | Pipistrellus pipistrellus | Monodnaviria | 5023581 | Zophobas morio black wasting virus |
| C05 | Care_KW_S21 | Pipistrellus pipistrellus | Monodnaviria | 21425 | Zophobas morio black wasting virus |
| W12 | Wild_Hirst Wood_S15 | Nyctalus noctula | Monodnaviria | 192 | Zophobas morio black wasting virus |
| W3 | Wild_DG_S3 | Myotis daubentonii | Monodnaviria | 9271 | Protoambidensovirus incertum4 |
| W3 | Wild_DG_S2 | Myotis daubentonii | Monodnaviria | 121 | Protoambidensovirus incertum4 |
| W3 | Wild_DG_S3 | Myotis daubentonii | Monodnaviria | 57 | Bee densovirus 2 |
| W3 | Wild_DG_S3 | Myotis daubentonii | Monodnaviria | 2690 | Bee densovirus 2 |
| W2 | Wild_DG_S2 | Myotis daubentonii | Monodnaviria | 238 | Bat parvovirus 3 |
| W2 | Wild_DG_S2 | Myotis daubentonii | Riboviria | 58 | NA |
| W2 | Wild_DG_S2 | Myotis daubentonii | Riboviria | 91 | bat chromosome |
